# Elucidating genes sufficient for viral entry into cells through sequential genome-wide CRISPR activation screens

**DOI:** 10.64898/2026.03.06.710083

**Authors:** Timothy Chai, Alicia Wong, Qingqing Yin, Isabel von Creytz, Jonathan S. Weissman, Reuben A. Saunders, Joseph B. Prescott, Kyle M. Loh

## Abstract

A preeminent goal of virology is to discover cellular genes that mediate virus entry. Genome-wide loss-of-function screens can illuminate single genes *necessary* for virus entry, but are stymied by genetic redundancy. Here we report a genome-wide CRISPR activation screening strategy to discover single genes that are *sufficient* for viral entry into normally-uninfectable cells. Sequential rounds of viral infection vastly enhanced screening sensitivity. This sequential screening strategy was generalizable to two unrelated viruses—Ebola and rabies viruses—and could broadly accelerate the discovery of viral entry factors.

## Introduction

Discovering the cellular proteins that mediate virus entry is imperative to therapeutically interdict viral entry into cells. Cellular entry of enveloped viruses encompasses multiple steps, including attachment to a host cell, followed by membrane fusion^1,2^. Historically, multiple approaches have been used to discover cellular proteins required for viral entry. First, if a viral envelope protein stably binds a cellular receptor, it can be biochemically discovered by immunoprecipitation, as exemplified by SARS coronavirus^3^, MERS coronavirus^4^, and Nipah virus^5^, among other viruses. However, immunoprecipitation requires stable and high-affinity binding of a viral envelope protein to a cellular protein. Second, genetic deletion represents another elegant approach to discover genes required for viral entry^6,7^, and has been used to elucidate cellular receptors of Lassa virus^8^, Ebola virus^9^, Rift Valley fever virus^10^, Lujo virus^11^, New World hantaviruses^12^, mouse norovirus^13^, and other viruses. While genetic deletion can prove successful if there is a single cellular gene *necessary* for viral entry, it can be stymied if there are multiple redundant entry factors such that loss of a single gene is compensated by the expression of other genes (**Fig. 1A**). For instance, human cytomegalovirus^14^ and Nipah virus^15^ can each exploit multiple receptors for cellular entry, thus challenging genetic screens that entail deletion of one gene at a time.

**Figure 1:**
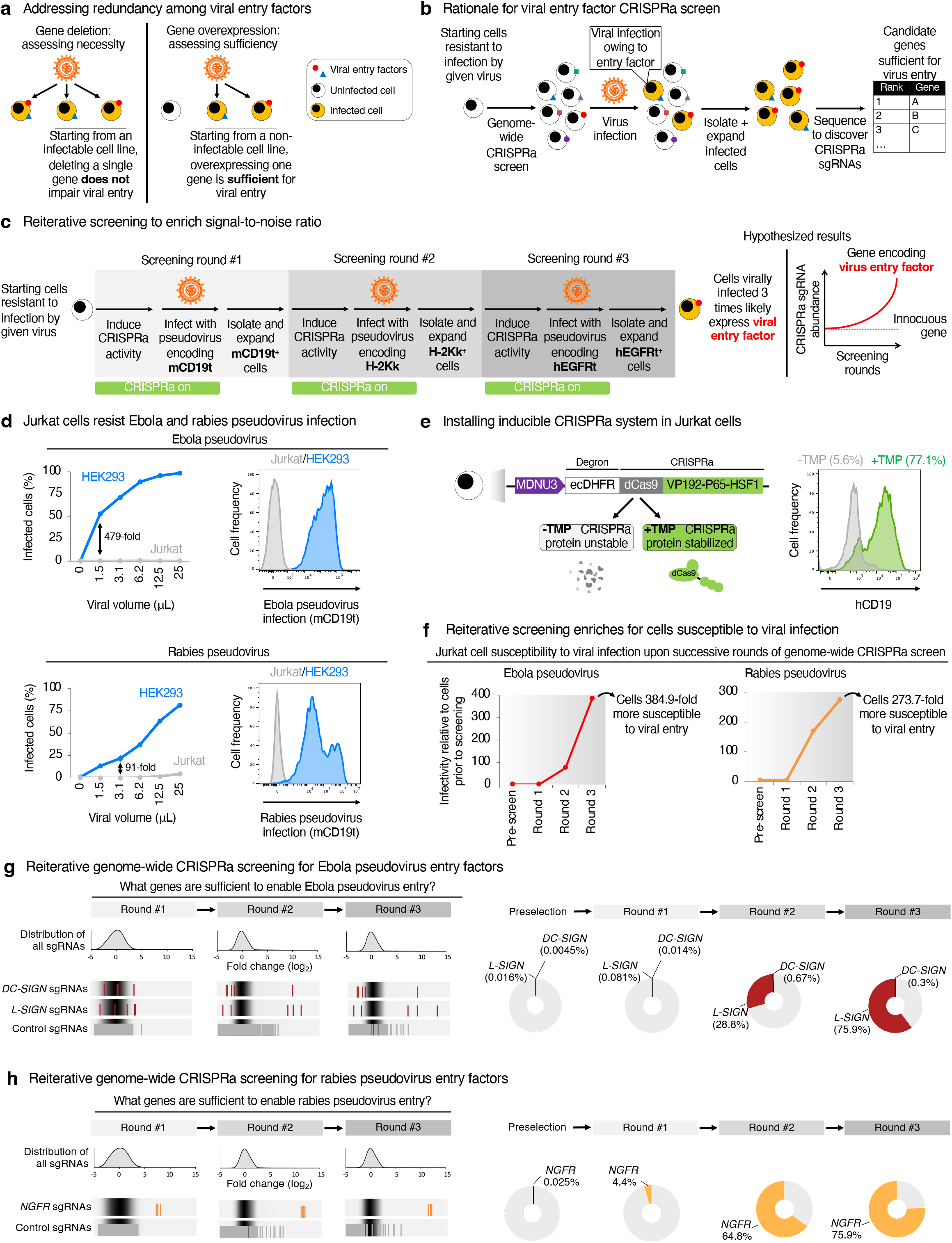
Reiterative genome-wide CRISPRa screens define cellular genes sufficient for virus entry. (A) Summary of gene deletion vs. gene overexpression approaches. (B) Rationale for the present study. (C) Experimental outline of the present study. (D) HEK293 and Jurkat cells were inoculated with different volumes of Ebola or rabies pseudovirus encoding a cell-surface marker (mCD19t; a truncated mutant of mouse CD19), followed by flow cytometry to determine the percentage of infected cells. This revealed that Jurkat cells are largely refractory to Ebola or rabies pseudovirus entry, relative to HEK293 cells. (E) Construction of a clonal Jurkat cell line, known as “Jurkat C6”, stably expressing a degron-tagged CRISPRa construct (*left*). Jurkat C6 cells were transduced with sgRNA targeting the endogenous human *CD19* gene, in the presence or absence of TMP (1 μM) for 3 days, followed by flow cytometry to detect hCD19 expression (*right*). (F) At each of the indicated points of the genome-wide CRISPRa screen, the cell population was challenged with Ebola or rabies pseudovirus, and cell infectivity was evaluated by flow cytometry. Upon successive rounds of the screen, the cell population became progressively more susceptible to infection to either Ebola or rabies pseudovirus, respectively. (G) sgRNA distribution upon successive rounds of the genome-wide CRISPRa screen for Ebola pseudovirus entry. (H) sgRNA distribution upon successive rounds of the genome-wide CRISPRa screen for rabies pseudovirus entry.

Importantly, gene overexpression enables the discovery of genes *sufficient* for virus entry (**Fig. 1A**). We were inspired by classical studies that employed gene overexpression to discover cellular entry factors for HIV-1^16^ and Nipah virus^17^; these studies measured cell-to-cell fusion mediated by viral envelope proteins. We expanded this gene overexpression strategy to the genome-wide scale using CRISPR activation (CRISPRa)^18,19^, incorporated sequential rounds of screening (as described below), and turned to a different experimental assay, namely virus entry.

Starting from a cell line that is refractory to virus infection, we performed genome-wide CRISPRa screens^18^ to test if overexpression of a single gene is *sufficient* for viral entry (**Fig. 1B**). One challenge with this approach is the signal-to-noise ratio. In a genome-wide CRISPRa library, only a small number of sgRNAs are expected to activate genes encoding *bona fide* viral entry factors. After a single round of infection by a virus encoding a selectable marker, successfully infected cells will comprise two subsets: cells harboring an sgRNA that activated a true entry factor (and that were infected *because* of this entry factor), and cells that were infected by chance. Critically, only the former population carries a heritable genetic element— the sgRNA—that will reproducibly enhance viral entry once more upon a subsequent round of viral infection. By contrast, cells infected by chance are no more likely than any other cell to be infected again upon another round of infection. We therefore reasoned that reiterative rounds of viral infection would progressively enrich for cells harboring sgRNAs targeting authentic entry factors, while diluting out stochastically infected cells (**Fig. 1C**). In each round, cells were virally infected, the small number of successfully infected cells were purified and expanded, and then subjected to an additional round of viral infection with a virus encoding a distinct selectable marker (**Fig. 1C**). We demonstrate that this serial genome-wide CRISPRa screening strategy can identify cellular entry factors for two unrelated viruses—Ebola virus and Rabies virus—thus suggesting its broad utility. More generally, this method of reiteratively exposing cells to a selective pressure to amplify rare heritable events may also benefit other genome-wide screening approaches beyond those that concern viral entry.

## Results

### Reiterative genome-wide CRISPRa screens for viral entry

To study viral entry, we generated pseudotyped lentiviruses bearing the envelope protein (glycoprotein) of either Ebola virus (Makona variant^20^) or rabies virus (CVS-N2C variant^21^) (**Fig. S1A**). Pseudotyped lentiviruses enable precise studies of viral entry, because they can enter target cells through the envelope protein displayed on their surface, but subsequently cannot replicate^22^. Attesting to the relevance of the viral variants studied here, Ebola virus (Makona variant) emerged in the 2013-2014 Guinea outbreak^23^, whereas rabies virus (CVS-N2C variant) is neurotropic *in vivo* and *in vitro*^24^.

Jurkat E6.1 cells—immortalized T cells—were selected for our screen, as they were up to 290-479 times less susceptible to infection by lentiviruses pseudotyped with either rabies or Ebola envelope proteins^25-27^, relative to HEK293T/17 cells (**Fig. 1D**). Given that Jurkat cells were still susceptible to infection by lentiviruses pseudotyped with vesicular stomatitis virus (VSV) envelope protein (**Fig. S1B**), this suggests that Jurkat cells selectively lack components necessary for Ebola and rabies entry. Jurkat cells thus provide an ideal negative background to test whether overexpression of a given gene is *sufficient* to enable Ebola or rabies entry. We engineered Jurkat cells with an inducible CRISPRa system, whereby the CRISPRa protein (dCas9-VP192-P65-HSF1)^19^ was fused to the DHFR degron domain^28^, such that the CRISPRa protein was stabilized upon treatment with a DHFR-binding small molecule (TMP)^28^ (**Fig. 1E, Fig. S1C**).

We then established a reiterative CRISPRa screening platform. We delivered a lentiviral genome-wide sgRNA library^29^ into CRISPRa-expressing Jurkat cells, followed by TMP treatment to activate the CRISPRa machinery and to achieve human gene overexpression. This pool of Jurkat cells was then exposed to a first round of infection by either Ebola pseudovirus or rabies pseudovirus, each encoding a selectable cell-surface marker (CD19) (**Fig. 1C**). We expected that most Jurkat cells should resist pseudovirus infection, except for rare Jurkat cells (1) in which CRISPRa had induced an entry factor sufficient for viral entry, or (2) that were infected by chance, given that Ebola and rabies pseudoviruses can enter wild-type Jurkat cells, albeit at very low frequency.

CD19-expressing cells (i.e., cells successfully infected by either Ebola or rabies pseudovirus in the first round) were isolated by magnetic-activated cell sorting (MACS) and expanded (**Fig. 1C**). They were then serially subjected to a second round of CRISPRa activation and viral infection by either Ebola or rabies pseudovirus encoding a different cell-surface marker (H-2Kk), and H2Kk-expressing cells were isolated and expanded once more (**Fig. 1C**). This was then followed by a third round of CRISPRa activation and infection by either Ebola or rabies pseudovirus encoding a distinct cell-surface marker (EGFR), followed by cell isolation and expansion. Jurkat cells expressing all three surface markers (e.g., CD19, H-2Kk, and EGFR) represent cells that had been successfully infected by pseudovirus on three separate occasions (**Fig. 1C**). We hypothesized that in such cells, it was probable that CRISPRa had induced a viral entry factor.

Upon each round of iterative enrichment, the cell population became progressively more susceptible to Ebola or rabies pseudovirus infection, consistent with CRISPRa-mediated upregulation of viral entry factors (**Fig. 1F**). Because stochastically infected cells carry no heritable advantage, they are unlikely to persist into subsequent rounds of selection, whereas cells wherein CRISPRa induced a true entry factor are likely to be progressively enriched in each round. By round three of the screen, the effects of progressive selection became pronounced: the cell population became 384.9-fold and 273.7-fold more susceptible to infection by Ebola pseudovirus or rabies pseudovirus, respectively (**Fig. 1F**). Genomic sequencing was performed after each screening round to track sgRNA enrichment over time (**Table S1**).

### Viral entry factors defined by reiterative genome-wide CRISPRa screening

In successive rounds of the Ebola pseudovirus screen, CRISPRa sgRNAs targeting *L-SIGN* (otherwise named *CLEC4M* or *CD299*) progressively increased in abundance: they comprised 0.016% of the sgRNA library prior to screening, 0.1% after round 1, 28.8% after round 2, and 75.9% after round 3 (**Fig. 1G, Table S1**). The Ebola pseudovirus screen also highlighted a related family member, *DC-SIGN* (otherwise named *CLEC4L* or *CD209*): *DC-SIGN* sgRNA abundance increased by two orders of magnitude through sequential screening rounds (**Fig. 1G, Table S1**). L-SIGN and DC-SIGN are transmembrane proteins that have been previously implicated as Ebola virus attachment factors^30-32^, as described further below. These dynamics illustrate how successive rounds of screening enhance the signal-to-noise ratio. Reiterative screening thus significantly enriches true-positive hits within genome-wide CRISPRa screens, particularly those with initially weak signals.

Consecutive rounds of rabies pseudovirus screening progressively enriched for CRISPRa sgRNAs targeting *NGFR*, which comprised of 0.025% of sgRNAs prior to screening, but rose to 4.5%, 65%, and 75% of total sgRNAs by rounds 1, 2 and 3, respectively (**Fig. 1H, Table S1**). NGFR (Nerve Growth Factor Receptor, alternately named p75) is a transmembrane protein previously implicated in rabies virus entry^33^ and physically interacts with rabies glycoprotein^34,35^. Taken together, reiterative screening rounds increase screening sensitivity and specificity in the context of both Ebola and rabies pseudovirus entry.

### Validation of genes sufficient for viral entry that emerged from reiterative genome-wide CRISPRa screens

We validated these genome-wide CRISPRa viral entry screens by overexpressing the candidate genes in Jurkat cells. While Jurkat cells were highly refractory to rabies pseudovirus infection, *NGFR* expression sufficed to enable rabies pseudovirus entry into Jurkat cells (**Fig. 2A, Fig. S2A**). This phenotype was observed after *NGFR* expression accomplished by either CRISPRa (to activate the endogenous promoter at physiological levels^36^) or an exogenous complementary DNA (cDNA) transgene (**Fig. 2A, Fig. S2A**). It was previously known that *NGFR* expression enables rabies virus infection of BSR cells^33^, and our study extends these findings to Jurkat cells. Collectively, this suggests that *NGFR* is sufficient for rabies virus entry across multiple cellular contexts.

**Figure 2:**
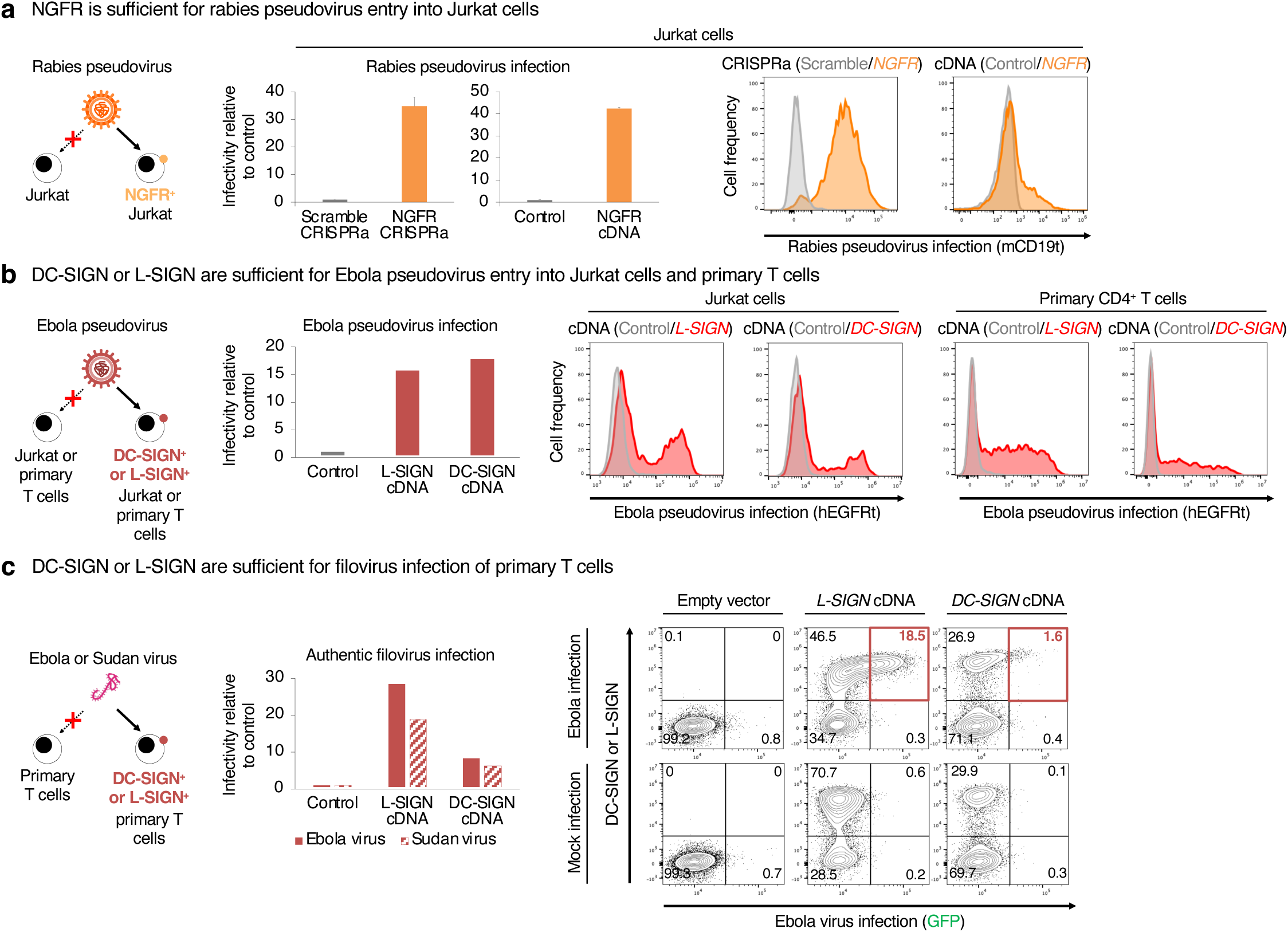
Validation of genome-wide CRISPR activation screening hits. A) *NGFR* was expressed in Jurkat C6 cells using CRISPRa, or alternatively, Jurkat cells using cDNA expression. *NGFR*-expressing or control cells were then inoculated with rabies pseudovirus encoding mCD19t. Flow cytometry was then performed to determine the percentage of infected cells. This revealed that *NGFR* expression significantly increased the susceptibility of Jurkat cells to rabies pseudovirus infection. B) *L-SIGN* or *DC-SIGN* were expressed in Jurkat or primary human CD4^+^ T cells using cDNA expression. *L-SIGN*-expressing, *DC-SIGN*-expressing, or control cells were inoculated with rabies pseudovirus encoding hEGFRt (a truncated mutant of human EGFR). Flow cytometry was then performed to determine the percentage of infected cells. This revealed that *L-SIGN* or *DC-SIGN* expression significantly increased the susceptibility of Jurkat cells and primary T cells to Ebola pseudovirus infection. C) *L-SIGN* or *DC-SIGN* were expressed in primary human CD4^+^ T cells using cDNA expression, and then *L-SIGN*-expressing, *DC-SIGN*-expressing, or control cells were inoculated with *GFP*-expressing Ebola virus^37^ or *zsGreen*-expressing Sudan virus^38^ under BSL4 containment. Flow cytometry was then performed to determine the percentage of infected cells. This revealed that *L-SIGN* or *DC-SIGN* expression significantly increased the susceptibility of primary T cells to authentic Ebola and Sudan virus infection.

While Jurkat cells were refractory to Ebola pseudovirus infection, expression of either *DC-SIGN* or *L-SIGN* was sufficient to enable Ebola pseudovirus entry, as observed upon either CRISPRa-or cDNA-mediated expression of these two genes (**Fig. 2B**). This is consistent with past findings that *DC-SIGN* or *L-SIGN* expression enhances Ebola pseudovirus infection of multiple immortalized cell lines^27,30-32^.

Expanding beyond immortalized cell lines, we turned to a more physiological context: human primary CD4^+^ T cells. While primary human CD4^+^ T cells were uninfectable by Ebola pseudovirus^32^, we discovered that forced expression of either *DC-SIGN* or *L-SIGN* was sufficient to enable Ebola pseudovirus entry into human primary CD4^+^ T cells (**Fig. 2B, Fig. S2B**). This contrasts with a prior report that *DC-SIGN* or *L-SIGN* expression in primary human CD4^+^ T cells did not enhance their susceptibility to Ebola pseudovirus^32^. The differences between our studies may be ascribed to the levels of gene overexpression: we found that T cells expressing the highest levels of *DC-SIGN* or *L-SIGN* were preferentially infected by Ebola pseudovirus (**Fig. S2B**).

Cognizant that pseudoviruses provide imperfect models of viral entry, we infected primary human CD4^+^ T cells with authentic Ebola virus under biosafety-level-4 (BSL4) containment. We employed a recombinant Ebola virus (Mayinga isolate) engineered with an additional *GFP* transcriptional unit^37^. Primary CD4^+^ T cells were highly resistant to Ebola infection, whereas expression of either *DC-SIGN* or *L-SIGN* enabled Ebola virus infection, as denoted by GFP fluorescence (**Fig. 2C**). We exploited the fact that engineered T cells displayed heterogeneous levels of DC-SIGN or L-SIGN expression to demonstrate that Ebola infection correlated with the detectable cell-surface expression of DC-SIGN or L-SIGN by T cells (**Fig. 2C**), thus associating these proteins with viral entry. Similarly, *DC-SIGN* or *L-SIGN* expression was sufficient to enable T cell infection by Sudan virus, another filovirus in genus *Ebolavirus* (**Fig. 2C**); for these experiments, we employed a recombinant Sudan virus (Gulu-808892 isolate) engineered with an additional *zsGreen* transcriptional unit^38^. Interestingly, despite initial Ebola and Sudan virus infection of primary CD4^+^ T cells, sustained viral replication was not observed (**Fig. S2C**), intimating that a T cell-intrinsic restriction factor may block filovirus genome replication.

Taken together, these results suggest that DC-SIGN or L-SIGN are sufficient for Ebola virus entry, and confirm that phenotypes observed in Jurkat cells are recapitulated in a physiologically relevant system.

## Discussion

Herein we present a genome-wide CRISPRa screening approach to discover genes whose expression is *sufficient* for viral entry into normally-uninfectable cells. This screening strategy holds promise for scenarios wherein a virus may exploit multiple redundant entry factors^14,15^. It thus complements genetic loss-of-function screens to discover genes that are *necessary* for viral entry but might be challenged by genetic redundancy.

Paramount to our approach is the use of reiterative rounds of viral infection to progressively enrich for cells harboring sgRNAs that confer viral susceptibility. The rationale for this approach rests on simple population genetics logic. After a single round of viral challenge in the context of a genome-wide CRISPRa library, the pool of successfully infected cells will inevitably contain a mixture of cells: “true positive” cells in which CRISPRa activated a gene encoding an authentic viral entry factor and “false positive” cells infected by chance. However, only true positives carry a heritable genetic element (the sgRNA) that reproducibly confers a selective advantage upon subsequent rounds of viral challenge. False positives, lacking any such heritable advantage, are selected against in each subsequent screening round. Over successive rounds, this differential selection exponentially enriches for true-positive sgRNAs while depleting stochastic contaminants, thereby transforming a weak initial signal into a dominant one. The dynamics observed in our screens are consistent with this model. For instance, in our Ebola pseudovirus entry screen, sgRNAs targeting *L-SIGN* rose from 0.016% of the library (prior to screening) to 75.9% (after three rounds), whereas initially enriched but biologically irrelevant sgRNAs such as those targeting *PLA2G4B* were depleted to near-undetectable levels by round 2 (**Fig. 1G, Table S1**).

Our iterative enrichment strategy contrasts with previous genetic screens that employed a continuous selective pressure—such as a toxic drug^39^ or a continuously replicating and lytic virus (e.g., SARS-CoV-2^40,41^, norovirus^13^, or pseudotyped VSV^8-12^)—to select mutant cells that can resist this selective pressure. In such cases, the virus continuously replicates until all susceptible cells are lysed, thus imposing *negative* selection^8-13,40,41^. Our screen instead requires *positive* selection of infected cells and thus a lytic virus is not fit for purpose. We thus reiteratively challenged cells with pseudoviruses encoding different cell-surface markers to progressively enrich for sgRNA-expressing cells susceptible to viral infection. Despite relatively modest signals for authentic viral entry factors in the first round of screening, they are substantially enriched upon consecutive rounds of viral infection (**Fig. 1G-H**), thus emphasizing the importance of reiterative screening. Two past studies applied CRISPRa screens to study SARS-CoV-2 entry factors, but only employed one round of screening^42,43^. The low signal-to-noise ratio afforded by one round of screening may contribute to why few hits emerging from such screens were subsequently validated as authentic SARS-CoV-2 entry factors^42,43^, although other factors—such as differences in cell lines, library coverage, and the biology of ACE2-mediated entry—may have also palyed a role.

Critically, we exploited a degron system to transiently activate CRISPRa for several day pulses prior to and during viral infection, but subsequently deactivated CRISPRa. In a genome-wide CRISPRa screen, CRISPRa will activate genes encoding cell-cycle activators (e.g., proto-oncogenes) that would otherwise skew the genetic composition of the cell population and obscure discovery of viral entry factors. Transient expression of CRISPRa also minimizes its cellular toxicity^44^.

As with any method, this reiterative genome-wide CRISPRa screening approach also harbors limitations. First, it can only discover entry factors that are limiting for viral entry in the specific cell-type being studied. Second, genes repressed by constitutive heterochromatin in each cell-type are less activatable by CRISPRa^36,45^. Both limitations may be potentially addressed by extending this genome-wide CRISPRa screening approach to multiple cell-types, wherein each cell-type presumably the limiting viral entry factor(s) and landscape of genes that can be activated by CRISPRa may differ. cDNA overexpression libraries could overcome these challenges; however they (1) are more challenging to work with, as variable cDNA lengths often lead to imbalanced library representation, and (2) can lead to super-physiological gene expression levels^46^. In any case, we demonstrate that our sequential genome-wide CRISPRa screening approach can highlight genes that are sufficient for viral entry across two unrelated viruses—Ebola and rabies viruses—thus suggesting broad applicability to diverse viruses. This approach may thus accelerate the discovery of viral entry factors, especially when redundant viral entry factors may prevail.

## Acknowledgements

We thank Markus Kainulainen for sharing recombinant Sudan virus and Owen Witte for sharing*VSV-G* plasmid. This study was supported by U.S. National Institutes of Health T32 T32GM007365 (T.C.), Stanford Medicine Dean’s Fellowship (Q.Y.), Robert Koch Institute intramural funds (J.B.P.), Amaranth Foundation (J.B.P., K.M.L.), and the Anonymous, Gilbert, and Weintz Families (K.M.L.). J.S.W. was supported as a Howard Hughes Medical Institute investigator. R.A.S. was supported as a Harvard Junior Fellow. K.M.L. was supported as a Packard Foundation Fellow, Human Frontier Science Program Young Investigator, Pew Scholar, Baxter Foundation Faculty Scholar, and the Anthony DiGenova Endowed Faculty Scholar.

## Supplemental Figure Legends

**Figure S1:**
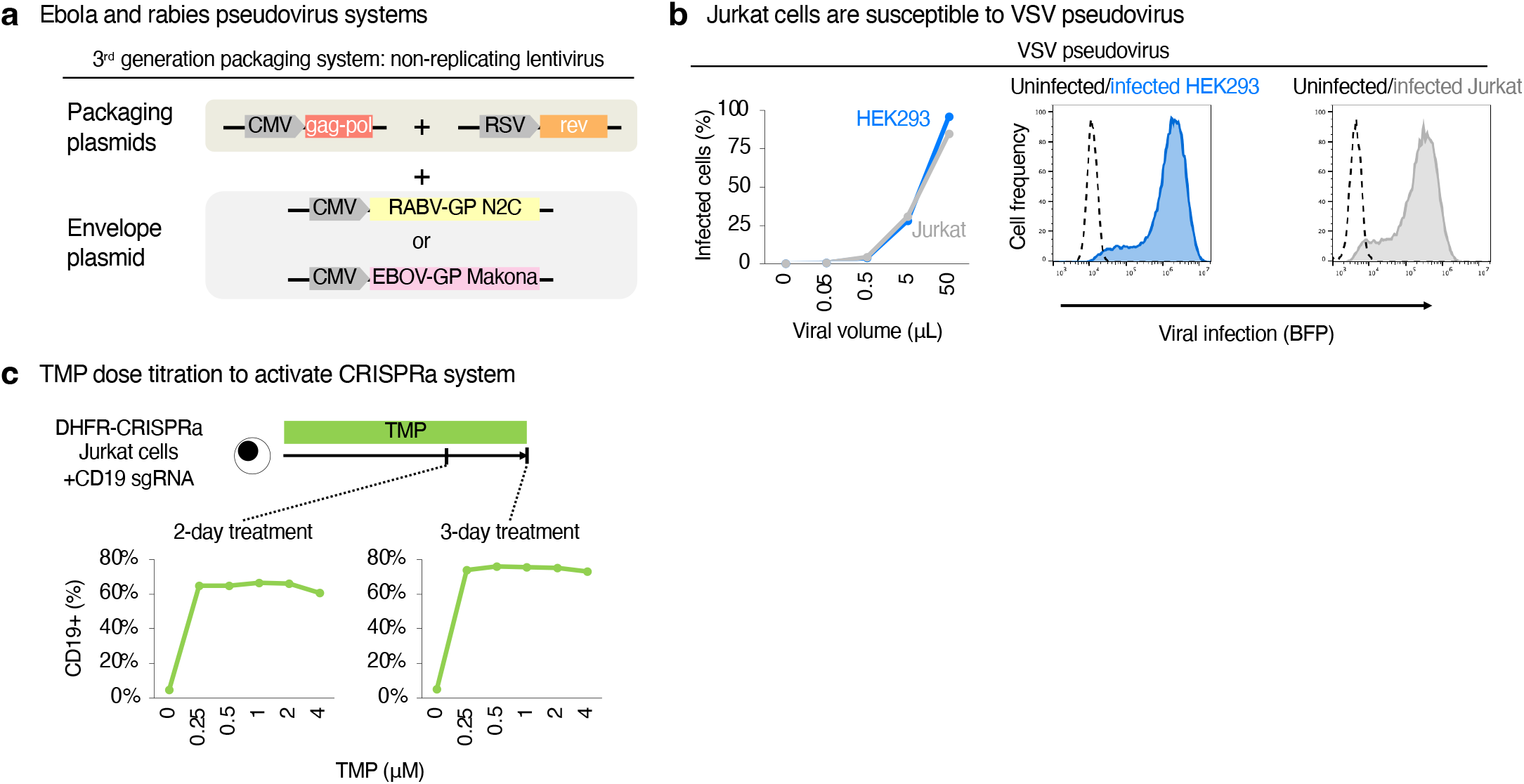
Establishing reiterative genome-wide CRISPRa screens for virus entry. A) Plasmids used to pseudotype non-replicating lentiviruses with either Ebola or rabies envelope proteins. EBOV-GP: glycoprotein of Ebola virus, Makona variant. RABV-GP N2C: glycoprotein of rabies virus, N2C variant. B) HEK293 and Jurkat cells were inoculated with different volumes of VSV envelope protein-pseudotyped lentivirus encoding a cell-surface marker (mCD19t; a truncated mutant of mouse CD19), followed by flow cytometry to determine the percentage of infected cells. This served as a positive control to confirm that Jurkat cells and HEK293 cells are both susceptible to VSV pseudovirus entry. C) Jurkat C6 cells were transduced with sgRNA targeting the endogenous human *CD19* gene, in the presence of different TMP concentrations (0-4 μM) for 2-3 days, followed by flow cytometry to detect human CD19 expression.

**Figure S2:**
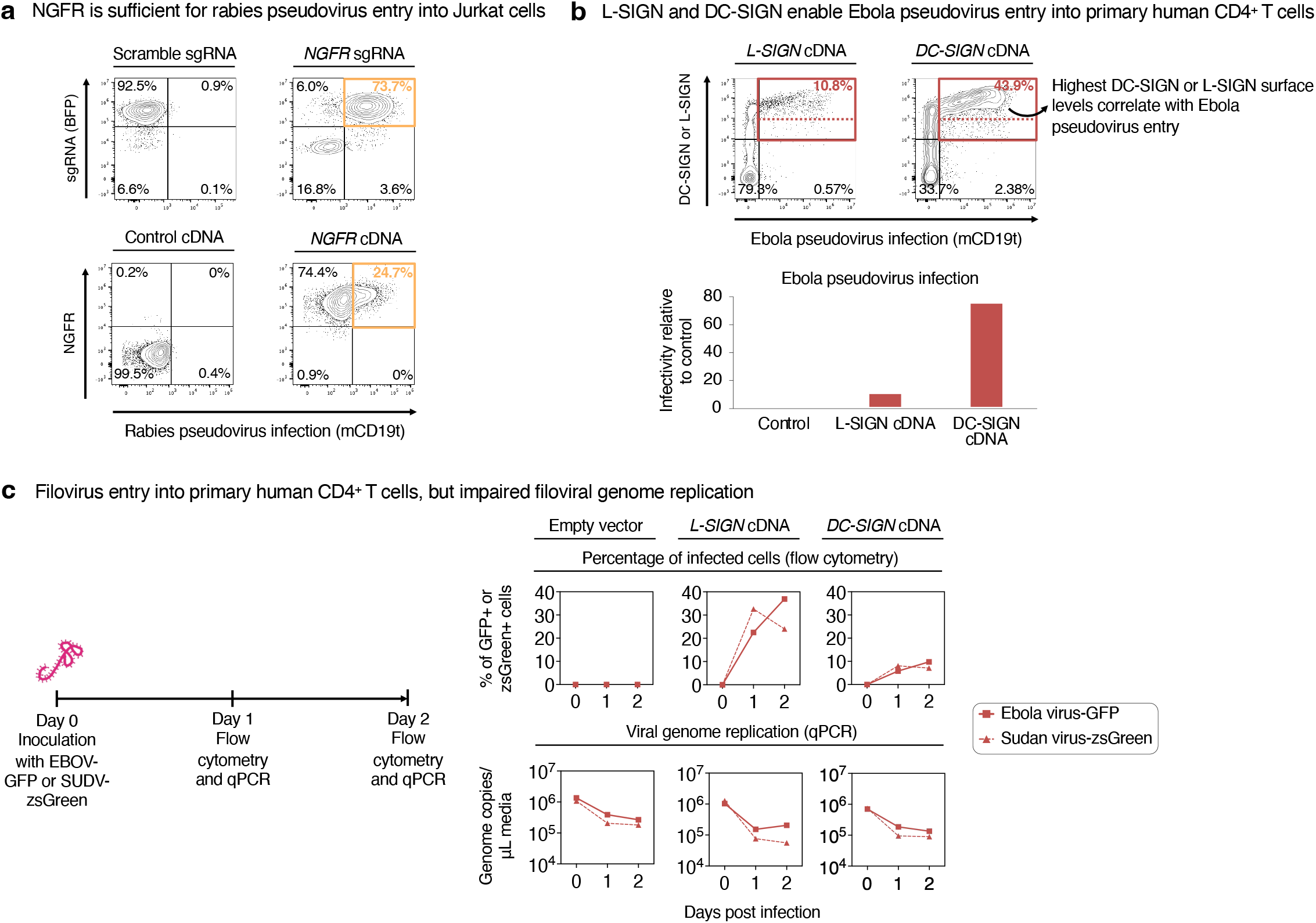
Validation of genome-wide CRISPR activation screening hits. A) *NGFR* was expressed in Jurkat C6 cells using CRISPRa, or alternatively, Jurkat cells using cDNA expression. *NGFR*-expressing or control cells were then inoculated with rabies pseudovirus encoding mCD19t. Flow cytometry was then performed to determine the percentage of infected cells. This revealed that *NGFR* expression significantly increased the susceptibility of Jurkat cells to rabies pseudovirus infection. As positive controls, flow cytometry was used to confirm successful delivery of the sgRNA construct as part of the CRISPRa workflow (as denoted by BFP expression) and that NGFR was expressed (upon cDNA expression). B) *L-SIGN* or *DC-SIGN* were expressed in primary human CD4^+^ T cells using cDNA expression. *L-SIGN*-expressing, *DC-SIGN*-expressing, or control cells were inoculated with rabies pseudovirus encoding hEGFRt (a truncated mutant of human EGFR). Flow cytometry was then performed to determine the percentage of infected cells. This revealed that *L-SIGN* or *DC-SIGN* expression significantly increased the susceptibility of Jurkat cells and primary T cells to Ebola pseudovirus infection. Cells expressing the highest levels of L-SIGN and DC-SIGN were preferentially infected by Ebola pseudovirus. C) *L-SIGN* or *DC-SIGN* were expressed in primary human CD4^+^ T cells using cDNA expression, and then *L-SIGN*-expressing, *DC-SIGN*-expressing, or control cells were inoculated with *GFP*-expressing Ebola virus^37^ or *zsGreen*-expressing Sudan virus^38^ under BSL4 containment. On days 0, 1, and 2 post-infection, flow cytometry was performed to determine the percentage of infected cells and qPCR was performed on cell culture supernatants to quantify viral genome replication. This revealed that *L-SIGN* or *DC-SIGN* expression enabled authentic Ebola and Sudan virus entry into primary human T cells, but viral genome replication was impaired, perhaps reflective of cell-intrinsic restriction factors.

## Supplemental Table Legends

**Table S1: Reiterative genome-wide CRISPRa screening results for Ebola and rabies pseudovirus entry** Each tab of the spreadsheet describes sgRNA representation in cell populations at a distinct phase of the reiterative genome-wide CRISPRa screen: prior to screening, and after rounds 1, 2, and 3 of the screen. sgRNAs represented in the cell population were grouped together if they targeted the same gene, and genes were rank ordered by the collective abundance of all their sgRNAs. Each line represents a distinct sgRNA.

## Methods

### Cell lines and culture

Jurkat E6.1 cells (ATCC, TIB-152) were cultured in RPMI 1640 (Thermo Fisher, 72-400-120), supplemented with 10% fetal bovine serum (FBS, Thermo Fisher, A5670801) and 1% penicillin-streptomycin (Thermo Fisher, 15-140-122).

HEK293T/17 cells (ATCC, CRL11268) were cultured in DMEM (Thermo Fisher, 10566024), supplemented with 10% fetal bovine serum (FBS, Thermo Fisher, A5670801) and 1% penicillin-streptomycin (Thermo Fisher, 15-140-122).

Primary human CD4^+^ T cells were isolated from healthy donor peripheral blood mononuclear cells (PBMCs) by negative selection using the EasySep Human CD4^+^ T Cell Isolation Kit (STEMCELL Technologies, 17952) according to the manufacturer’s instructions. For activation and expansion, CD4^+^ T cells were cultured in AIM-V medium (Thermo Fisher, 12055083) supplemented with 5% heat-inactivated human serum (Millipore Sigma, H3667-100ML), 1× GlutaMAX (Thermo Fisher, 35050061), 55 nM β-mercaptoethanol (Thermo Fisher, 21985023), 5 ng/mL recombinant human IL-7 (R&D Systems, 207-IL-050), and 0.5 ng/mL recombinant human IL-15 (PeproTech, 200-15). Cells were activated immediately after isolation using Human T-Activator CD3/CD28 beads (Thermo Fisher, 11132D) at a 1:1 bead-to-cell ratio. Beads were removed prior to experimentation using a magnet (Thermo Fisher, 12302D) according to the manufacturer’s instructions. All primary human cell experiments were approved by the Stanford Blood Center under an institutional review board-approved protocol and informed consent was obtained from all donors.

All cells in this study were cultured at 37°C in a humidified atmosphere with 5% CO_2_.

### Generation of degron-tagged CRISPRa Jurkat cells

In brief, we engineered a clonal Jurkat E6.1 line expressing a degron-tagged CRISPRa protein (DHFR-dCas9-VP192-P65-HSF1)^19^; we refer to this clonal cell line as “Jurkat C9” throughout this study. The degron tag (*E. coli* DHFR) constitutively destabilizes the protein-of-interest, in this case the CRISPRa protein^28^. However, addition of the small molecule trimethoprim (TMP) stabilizes the CRISPRa protein^28^.

Specifically, Jurkat E6.1 cells were transduced with a lentiviral vector encoding the *MNDU3* promoter driving the expression of DHFR-dCas9-VP192-P65-HSF1-T2A-hNGFRt. hNGFRt refers to a truncated human NGFR/CD271 protein (referred to as “NGFRt”), a cell-surface marker that enables the facile selection of transduced cells.

After lentiviral transduction, Jurkat cells were stained with a NGFR/CD271 PE antibody (BioLegend, 345106), and single NGFR^+^ cells were isolated and deposited into individual wells of a 96-well plate using fluorescence-activated cell sorting (FACS) to generate clonal cell lines. Individual clonal cell lines were expanded and screened for TMP-dependent CRISPRa activity by introducing an sgRNA targeting the endogenous *CD19* locus and quantifying CD19 surface induction by flow cytometry using a CD19 PE antibody (BioLegend, 302254), in the presence or absence of 1 μM TMP (Sigma-Aldrich, T7883-5G). The clonal CRISPRa-expressing cell line exhibiting the best TMP-inducible CRISPRa activity was designated “Jurkat C9”, and was used for subsequent experimentation.

### Selection of viral envelope proteins to generate pseudotyped lentiviruses

To produce pseudotyped lentiviral particles, we utilized plasmids encoding the viral envelope glycoproteins from the following viruses:

- **Ebola virus, Makona variant, C15 isolate** (H.sapiens-wt/GIN/2014/Makona-Kissidougou-C15): The genomic sequence of this viral isolate was determined by Baize et al., 2014 from an Ebola patient in the 2013-2014 Guinea outbreak of Ebola virus disease^23^. The genomic sequence of this viral isolate has been deposited to NCBI (accession number KJ660346.2). This study employed a plasmid encoding the glycoprotein of this viral isolate (EBOV-GP), which was generated by Diehl et al., 2016^20^ and deposited on Addgene (86021). Ebola virus is a member of the species *Orthoebolavirus zairense*; genus *Orthoebolavirus*; family *Filoviridae*; order *Mononegavirales*^47^.
- **Rabies virus, CVS-24 strain, N2C variant**: The challenge virus standard (CVS) strain of rabies virus was derived from the original Pasteur strain in 1882^48^. The CVS strain was employed by the U.S. National Institutes of Health (NIH) to determine rabies vaccine potency, and was also was used to manufacture rabies vaccine in the United States^48^. The CVS-24 strain has been exclusively passaged in mouse brains^24^. The N2C variant of the CVS-24 strain exhibits neurotropism *in vivo* and *in vitro*^24^. This study employed a plasmid encoding the glycoprotein of the N2C viral variant (RABV-G), which was generated by Mentis et al., 2006^49^ and deposited on Addgene (19712). Rabies virus is a member of the species *Lyssavirus rabies*; genus *Lyssavirus*; family *Rhabdoviridae*; order *Mononegavirales*.
- **Vesicular stomatitis virus**: To generate control pseudoviruses, we utilized a plasmid encoding the vesicular stomatitis virus envelope protein (VSV-G), provided by Dr. Owen Witte’s laboratory (University of California, Los Angeles)^50^. Vesicular stomatitis virus is a member of the species *Indiana vesiculovirus*; genus *Vesiculovirus*; family *Rhabdoviridae*; order *Mononegavirales*.

Both the Ebola and Rabies glycoprotein sequences were sub-cloned into a pcDNA3.3 expression backbone (Addgene, 172320) to drive robust viral envelope protein expression.

To generate the pseudotyped lentiviruses used in the reiterative genome-wide CRISPRa screen, we constructed three separate pCCL lentiviral vectors, each encoding a different cell-surface marker. First, pCCL-zsGreen-T2A-mCD19t, which encodes truncated mouse CD19 (mCD19t) and was employed in screening round 1. Second, pCCL-H-2Kkt, which encodes truncated mouse H-2Kk and was employed in screening round 2. Third, pCCL-hEGFRt, which encodes truncated human EGFR (hEGFRt) and was employed in screening round 3. Each cell-surface marker was chosen for compatibility with commercially available magnetic bead-based selection and antibodies.

### Generation of pseudotyped lentiviruses

In this study, we employed a 3^rd^ generation lentiviral packaging system to pseudotype lentiviral particles with a viral envelope protein of interest^51^, as described below. These lentiviral particles are self-inactivating, and upon infection of a target cell, cannot subsequently generate new infectious viral particles^51^.

Pseudotyped lentiviruses were produced in HEK293T/17 cells, as previously described^52^. Briefly, cells were seeded on plates coated with 0.01% poly-L-Lysine (R&D Systems, 3438-200-01) and cultured until they reached 80-90% confluency.

HEK293T/17 cells were then simultaneously transfected with four plasmids: (1) a pCCL-based lentiviral transfer plasmid, (2) the pMDL helper plasmid^51^, (3) the pREV helper plasmid^51^, and (4) a plasmid encoding a viral envelope protein (either EBOV-GP, RABV-N2C-G, or VSV-G, as described above). Transfection was performed using FuGENE HD (Promega, E2312) in Opti-MEM I Reduced Serum Medium (Thermo Fisher, 31985062). Following a 15-minute room temperature incubation, the transfection complex was added dropwise to the packaging cells. Two separate procedures were then used, depending on whether pseudotyped lentiviruses were being used to transduce cell lines or primary human T cells:

- **For future transduction of cell lines**: 16-20 hours post-transfection, the transfection medium was replaced with DMEM supplemented with 10% FBS. Viral supernatants were harvested 24 hours later, clarified by centrifugation, filtered through a 0.45 μm PES syringe filter (Millipore Sigma, SLHP033RS), and stored in single-use aliquots at -80°C.
- **For future transduction of primary human T cells**: 16-20 hours post-transfection, the transfection medium was replaced with AIM-V medium lacking serum. Lentiviral supernatants were then concentrated using 100 kDa Amicon ultra centrifugal filters (Millipore, UFC810008).

### Determination of cell line susceptibility to pseudotyped lentiviruses

HEK293T/17 and Jurkat C9 cell lines were infected with serial dilutions of lentiviruses that were pseudotyped with either EBOV-GP, RABV-G, or VSV-G. Cells were seeded in 96-well plates at 3×10^4^ cells per well in 50 μL of complete culture medium. Pseudovirus stocks were serially diluted twofold in complete medium to generate the following pseudoviral volumes per well: 25 μL, 12.5 μL, 6.25 μL, 3.125 μL, 1.562 μL, and 0 μL (mock control). All dilutions were performed in duplicate. Cells were infected with pseudotyped lentiviruses, and 72 hours later, were harvested and stained with antibodies against the relevant surface marker: mouse CD19 (Miltenyi Biotec, 130-111-884), mouse H2Kk (Miltenyi Biotec, 130-117-235), or human EGFR (R&D Systems, FAB9577R-100), and viability staining was performed using DAPI (Thermo Fisher Scientific, D1306). The percentage of marker-positive cells was quantified by flow cytometry.

The percentage of infected cells was plotted against the input viral volume (μL) to generate viral infectivity curves. Viral titers were calculated from the linear range of each curve. In subsequent experiments, these viral titers were used to determine the volume of viral stock required to achieve the desired multiplicity of infection (MOI) for each target cell line.

### Reiterative genome-wide CRISPRa screening to define viral entry factors

The human genome-wide CRISPRa sgRNA library (hCRISPRa-v2 [Addgene, 1000000091]) comprises 209,080 sgRNAs targeting 18,915 human genes (approximately 10 sgRNAs per gene), in addition to 3,790 non-targeting control sgRNAs^29^. The lentiviral sgRNA plasmid encodes a *U6* promoter driving sgRNA expression, followed by an *EF1A* promoter driving the expression of a *PuroR*-*T2A*-*BFP* cassette. This pooled CRISPRa sgRNA library was amplified and packaged into a VSV-G-pseudotyped lentiviral pool, as described above.

For the screen, 3×10^8^ Jurkat C9 cells were transduced with the pooled lentiviral CRISPRa sgRNA library at a multiplicity of infection (MOI) of 0.3. In these conditions, theoretically most infected cells (∼85.7%) receive a single sgRNA, thereby maintaining library representation of >500 cells per sgRNA. Transduction was performed in 350 mL total volume distributed across T225 flasks. Two days post-transduction, cells were selected with 2 μg/mL puromycin (Sigma-Aldrich, P8833-25MG) for 4 days to eliminate non-transduced cells.

Following puromycin selection, the CRISPRa protein was stabilized through addition of 1 μM TMP to the culture media for 72 hours prior to viral inoculation.

- **Screening round 1**: Jurkat C9 cells carrying the genome-wide CRISPRa sgRNA pool were treated with 1 μM TMP for 3 days (to stabilize the CRISPRa protein), and then inoculated with either Ebola or Rabies pseudovirus encoding mouse CD19t as a selectable cell-surface marker. The volume of Ebola or Rabies pseudovirus was chosen to achieve <1% infection in parental Jurkat cells, as determined from the aforementioned titration experiments. After 72 hours of pseudovirus inoculation, cells were harvested and mouse CD19t^+^ cells were enriched by magnetic-activated cell sorting (MACS) using anti-CD19 MicroBeads (Miltenyi Biotec, 130-121-301) according to the manufacturer’s instructions. The enriched population was expanded in culture for 5 days in media lacking TMP, until 3×10^6^ cells were obtained. A small aliquot was collected for flow cytometry to assess post-MACS enrichment purity. Both pre- and post-MACS samples were stained with anti-mouse CD19 antibody (Miltenyi Biotec, 130-111-884) and analyzed by flow cytometry to quantify enrichment efficiency. Two separate aliquots of 1×10^6^ cells each were cryopreserved in Bambanker (Bulldog Bio, BB05) for subsequent sgRNA sequencing. As a negative control, 2×10^8^ cells prior to MACS enrichment were also cryopreserved for subsequent sgRNA sequencing; these represented the “pre-enrichment” cell population.
- **Screening round 2:** 1×10^6^ Jurkat C9 cells from screening round 1 were treated with TMP for 3 days (to stabilize the CRISPRa protein), and then transduced with Ebola or Rabies pseudovirus encoding mouse H2Kk as a selectable cell-surface marker. After 72 hours of pseudovirus inoculation, cells were harvested and H2Kkt^+^ cells were enriched by MACS using MACSelect H2Kk beads (Miltenyi Biotec, 130-070-201) according to the manufacturer’s instructions. Enriched cells were expanded in culture for 11 days to obtain 2.5×10^6^ cells, which were then stained with an H-2kk antibody (Miltenyi Biotec, 130-117-235) to quantify enrichment efficiency, as per above. Two separate aliquots of 1×10^6^ cells each were cryopreserved for subsequent sgRNA sequencing.
- **Screening round 3**: 1×10^6^ Jurkat C9 cells from screening round 2 were treated with TMP for 3 days (to stabilize the CRISPRa protein), and then transduced with Ebola or Rabies pseudovirus encoding human EGFRt as a selectable cell-surface marker. After days of pseudovirus inoculation, hEGFRt^+^ cells were isolated by fluorescence-activated cell sorting (FACS) using an EGFR Alexa Fluor 647 antibody (R&D Systems, FAB9577R-100) on a BD FACSAria II FACS machine. Sorted cells were expanded for 10 days and cryopreserved.

### Genomic DNA isolation and sgRNA sequencing

The Nucleospin Blood Kit (Thermo Fisher, NC0389523) was used to extract genomic DNA (gDNA) from the Jurkat C9 cell population at four timepoints throughout the course of the genome-wide CRISPRa screen: prior to selection and the enriched populations from screening rounds 1, 2, and 3. sgRNA sequences were amplified by PCR and subjected to next-generation 150bp pair-end sequencing on an Illumina NextSeq 500 machine to quantify sgRNA abundance.

A custom Python script was used to extract and count sgRNA sequences from FASTQ files. Briefly, reads were filtered for the presence of upstream (5’ CCTTGTTG 3’) and downstream (5’ GTTTAAG 3’) flanking sequences that define the sgRNA cassette. For read 2 files (containing “R2” or “_2” in the filename), sequences were reverse complemented to ensure consistent orientation. The 20-nt sequence between the flanking regions represented the sgRNA. Reads containing the expected flanking sequences and yielding a 20-nucleotide-length sgRNA sequence were considered successfully identified. Extracted sgRNA sequences were aligned to the reference CRISPRa library (hCRISPRa-v2)^29^. For each sample, the number of reads mapping to each sgRNA was counted. Gene-level enrichment scores were calculated as the geometric mean of normalized counts (reads per million [RPM]) for all sgRNAs targeting that gene. Non-targeting control sgRNAs were used to estimate background noise and were adjusted by dividing their total counts by the number of control sgRNAs (3,789) and scaling to a reference value of 10 for comparative visualization.

### Validation of CRISPRa screening results

Genes that emerged from the genome-wide CRISPRa screen were validated in two ways: CRISPRa-mediated overexpression and cDNA-mediated overexpression.

To achieve CRISPRa-mediated overexpression, sgRNA sequences were cloned into the lentiviral sgRNA library backbone plasmid (Addgene, 84832) using the restriction sites BstX1 and BlpI. Separately, a non-targeting sgRNA sequence was also cloned as a negative control. This lentiviral sgRNA plasmid was then packaged into VSV-G-pseudotyped lentivirus, as described above. These sgRNA-carrying lentiviruses were then used to transduce CRISPRa-expressing Jurkat C9 cells. 2 days after lentiviral transduction, 2 μg/mL of puromycin was added to select for sgRNA-transduced cells. After 3 days of puromycin treatment, cells were treated with 1 μM TMP for 3 days to the stabilize CRISPRa protein and to activate the gene-of-interest. Cells were then transduced with either Ebola or Rabies pseudovirus, and 3 days later flow cytometry was performed to assess pseudovirus transduction (i.e., whether CRISPRa-mediated gene activation led to increased pseudovirus transduction).

To achieve cDNA-mediated overexpression, gBlocks encoding either *NGFR, L-SIGN*, or *DC-SIGN* were first synthesized by IDT. These gBlocks were then inserted into a dual promoter lentiviral vector with the *MNDU3* promoter driving expression of the gene-of-interest (“gene X”), followed by a *PGK* promoter driving hEGFRt as a selectable cell-surface marker (pCCL-MNDU3-gene X-PGK-hEGFRt). This lentiviral plasmid was packaged into a VSV-G pseudotyped lentivirus, as described above. Jurkat E6.1 cells or primary human CD4^+^ T cells were transduced with lentiviral vectors encoding either *NGFR, L-SIGN*, or *DC-SIGN*, or an empty vector as a negative control. Successful lentiviral transduction was confirmed by EGFR flow cytometry. Subsequently, cells were then transduced with either Ebola or Rabies pseudovirus, and 3 days later flow cytometry was performed to assess pseudovirus transduction (i.e., whether CRISPRa-mediated gene activation led to increased pseudovirus transduction).

### Preparation of concentrated filovirus stocks

Ebola and Sudan viruses are classified under Risk Group 4^53^. Consequently, all experiments with authentic Ebola and Sudan viruses were conducted under maximum containment conditions in the biosafety level 4 (BSL4) laboratory of the Robert Koch Institute, in accordance with standard operating procedures institutionally approved by the Robert Koch Institute. The following viruses were used:

- **Recombinant GFP-expressing Ebola virus, Yambuku variant, Mayinga isolate** (Ebola virus/Human-recombinant/COD/1976/Yambuku-Mayinga-GFP)^37^ was engineered by inserting a *GFP* cassette between the *NP* and *VP35* genes of Ebola virus, Yambuku variant, Mayinga isolate (Ebola virus/Human/COD/1976/Yambuku-Mayinga; genomic sequence reported in NCBI accession number AF086833.2). Recombinant GFP-expressing Ebola virus was rescued from Vero cells transfected with respective viral plasmids^37^, and was subsequently passaged on Vero E6 cells. Ebola virus is a member of the species *Orthoebolavirus zairense*; genus *Orthoebolavirus*; family *Filoviridae*; order *Mononegavirales*^47^. Recombinant Ebola virus was provided by the NIH, National Institute of Allergy and Infectious Diseases, Rocky Mountain Laboratories.
- **Recombinant zsGreen-expressing Sudan virus, Gulu-808892 isolate** (Sudan virus/Human-recombinant/UGA/2000/Gulu-808892-zsGreen)^38^ was engineered by inserting a *zsGreen-P2A* cassette immediately upstream of the *VP40* gene of Sudan virus, Gulu-808892 isolate (Sudan virus/Human/UGA/2000/Gulu-808892; genomic sequence reported in NCBI accession number KR063670.1). Recombinant zsGreen-expressing Sudan virus was rescued from HuH7 cells transfected with respective viral plasmids^38^, and was subsequently passaged on Vero E6 cells. Sudan virus is a member of the species *Orthoebolavirus sudanense*; genus *Orthoebolavirus*; family *Filoviridae*; order *Mononegavirales*. Recombinant Sudan virus was provided by the U.S. Centers for Disease Control and Prevention (CDC), Viral Special Pathogens Branch.

Highly concentrated stocks of GFP-expressing Ebola and zsGreen-expressing Sudan viruses were prepared prior to using them for infection studies. 270 mL of virus stock solution was concentrated by multiple rounds of centrifugation at 3000 rpm using Amicon Ultra-15 Centrifugal Filter Units (Merck, UFC9100), followed by a wash with a tenfold excess volume of DMEM, and finally the addition of FBS to a final concentration of 10%.

### Infection of primary T cells with filoviruses

Cryopreserved primary T cells lentivirally transduced with either an empty vector (negative control), an *L-SIGN* expression vector, or a *DC-SIGN* expression vector were thawed and resuspended in T cell medium containing a CD28 agonist antibody (2 μg/mL) and incubated in 6-well plates coated with a CD3 agonist antibody (1 μg/mL) for 3 days. Additional T cell medium was added at a 1:1 ratio on day 2 and day 3. On the third day, cells were transferred to new 6-well plates and on day 4, the cells were washed and resuspended in medium, in the absence of CD28 antibodies. 10^6^ T cells were then seeded per well of a 6-well plate. On the fifth day, cells were resuspended, and viable cells were using Trypan Blue. Subsequently, cells were transferred to 50 mL conical tubes and centrifuged at 500g for 5 minutes.

The medium was aspirated and 2.5×10^5^ cells were resuspended in 150 μL of medium containing either (1) GFP-expressing Ebola virus, at a multiplicity of infection (MOI) of 20, (2) zsGreen-expressing Sudan virus, at a multiplicity of infection (MOI) of 20, or (3) T cell medium (mock control). Three technical replicates were performed per condition. T cells were incubated in the viral inoculum for 1 hour at 37 °C with periodic agitation, and then centrifuged as above. The inoculum was then aspirated, and cells were resuspended in 1 mL of T cell medium and seeded at 2.5×10^5^ cells per well of a 12-well plate (0.5 mL of cell suspension/well) and incubated at 37 °C. Samples for flow cytometry and qPCR were harvested as described below.

### Flow cytometry analysis of filovirus-infected cells

1 or 2 days post-infection, T cells were harvested for flow cytometry. 280 μL of media laden with T cells was transferred to 96-well V-bottom plates and centrifuged at 600g for 3 minutes at 4 °C. The supernatant was aspirated, and then cells were resuspended in 45 μL of blocking buffer (Human TrueStain FcX^™^, diluted 1:200 [Biolegend, 422302]) for 10 minutes at room temperature.

Cells were then centrifuged and resuspended in 45 μL of buffer containing Live/Dead Yellow Stain (diluted 1:200 [Thermo Fisher, L34959]) and antibodies (all diluted 1:200; CD4 PE-Cy7, [Thermo Fisher, 25-0049-42], CD8A APC-Cy7 [Thermo Fisher, 47-0087-42], and DC-SIGN/L-SIGN APC [Thermo Fisher, 17-2999-42]) and incubated for 20 minutes in the dark. Cells were then washed in FACS wash buffer (PBS + 1% BSA + 0.1% NaN_3_) and centrifuged again. Subsequently, cells were resuspended and incubated in 10% formaldehyde overnight to inactivate materials for transfer out of the BSL4 laboratory. Afterwards, cells were centrifuged at 1000g for 10 minutes, resuspended in FACS buffer, and flow cytometric measurements were performed on a CytoFLEX S flow cytometry (Beckman Coulter). Flow cytometry data were analyzed using FlowJo v10 software (BD Life Sciences), using single color and fluorescence-minus-one controls for compensation, and isotype controls for thresholding.

### Quantitative PCR analysis of filovirus-infected cells

0, 1 or 2 days post-infection, T cells were harvested for viral RNA quantitation. 125 μL of media laden with T cells was placed in 2 mL tubes, and centrifuged at 600g for 5 minutes. The supernatant was then discarded and the cell pellet was resuspended in 600 μL of RLT buffer (Qiagen RNeasy Mini Kit [Qiagen, 74106]) containing 1% β-mercaptoethanol, in addition to 600 μL of 70% ethanol, to inactivate materials for transfer out of the BSL4 laboratory. Afterwards, RNA was extracted following the manufacturer’s instructions. Subsequently, one-step reverse transcription and quantitative PCR (qPCR) was performed using primers and FAM-labeled probes directed against the viral genome and the AgPath-ID One-Step RT-PCR Kit (Thermo Fisher, 4387391). The following PCR conditions were used: reverse transcription was performed at 45 °C for 15 minutes, followed by initial cDNA denaturation at 95°C for 10 minutes, followed by 43 cycles with denaturation at 95°C for 15 seconds, and annealing at 60°C for 1 minute.

To detect Ebola virus RNA, the following primer sequences were used:

- Forward primer: GTTCGTCTCCATCCTCTTGCA
- Reverse primer: TGAGGGAAAAGACCATGCTCA
- Probe: TGCTCCTTTCGCCCGACTTTTGAACC 6Fam-BBQ

To detect Sudan virus RNA, the following primer sequences were used:

- Forward primer: GGTGGTGTTGTTGACCCGTA
- Reverse primer: CATCGTCGTCGTCCAAATTGA
- Probe: TGAAGGCACCACAGGAGATCTTGATCT 6Fam-BBQ

## References

1 Marsh, M. & Helenius, A. Virus entry: open sesame. Cell 124, 729–740, doi:10.1016/j.cell.2006.02.007 (2006).

2 Jae, L. T. & Brummelkamp, T. R. Emerging intracellular receptors for hemorrhagic fever viruses. Trends Microbiol 23, 392–400, doi:10.1016/j.tim.2015.04.006 (2015).

3 Li, W. et al. Angiotensin-converting enzyme 2 is a functional receptor for the SARS coronavirus. Nature 426, 450–454, doi:10.1038/nature02145 (2003).

4 Raj, V. S. et al. Dipeptidyl peptidase 4 is a functional receptor for the emerging human coronavirus-EMC. Nature 495, 251–254, doi:10.1038/nature12005 (2013).

5 Negrete, O. A. et al. EphrinB2 is the entry receptor for Nipah virus, an emergent deadly paramyxovirus. Nature 436, 401–405, doi:10.1038/nature03838 (2005).

6 See, W. R., Yousefi, M. & Ooi, Y. S. A review of virus host factor discovery using CRISPR screening. mBio 15, e0320523, doi:10.1128/mbio.03205-23 (2024).

7 McDougall, W. M., Perreira, J. M., Reynolds, E. C. & Brass, A. L. CRISPR genetic screens to discover host-virus interactions. Curr Opin Virol 29, 87–100, doi:10.1016/j.coviro.2018.03.007 (2018).

8 Jae, L. T. et al. Virus entry. Lassa virus entry requires a trigger-induced receptor switch. Science 344, 1506–1510, doi:10.1126/science.1252480 (2014).

9 Carette, J. E. et al. Ebola virus entry requires the cholesterol transporter Niemann– Pick C1. Nature 477, 340–343, doi:10.1038/nature10348 (2011).

10 Ganaie, S. S. et al. Lrp1 is a host entry factor for Rift Valley fever virus. Cell 184, 5163–5178 e5124, doi:10.1016/j.cell.2021.09.001 (2021).

11 Raaben, M. et al. NRP2 and CD63 Are Host Factors for Lujo Virus Cell Entry. Cell host & microbe 22, 688–696.e685, doi:10.1016/j.chom.2017.10.002 (2017).

12 Jangra, R. K. et al. Protocadherin-1 is essential for cell entry by New World hantaviruses. Nature 563, 559–563, doi:10.1038/s41586-018-0702-1 (2018).

13 Orchard, R. C. et al. Discovery of a proteinaceous cellular receptor for a norovirus. Science 353, 933–936, doi:10.1126/science.aaf1220 (2016).

14 Martinez-Martin, N. et al. An Unbiased Screen for Human Cytomegalovirus Identifies Neuropilin-2 as a Central Viral Receptor. Cell 174, 1158–1171 e1119, doi:10.1016/j.cell.2018.06.028 (2018).

15 Negrete, O. A., Chu, D., Aguilar, H. C. & Lee, B. Single amino acid changes in the Nipah and Hendra virus attachment glycoproteins distinguish ephrinB2 from ephrinB3 usage. Journal of Virology 81, 10804–10814, doi:10.1128/JVI.00999-07 (2007).

16 Feng, Y., Broder, C. C., Kennedy, P. E. & Berger, E. A. HIV-1 entry cofactor: functional cDNA cloning of a seven-transmembrane, G protein-coupled receptor. Science 272, 872–877, doi:10.1126/science.272.5263.872 (1996).

17 Bonaparte, M. I. et al. Ephrin-B2 ligand is a functional receptor for Hendra virus and Nipah virus. Proceedings of the National Academy of Sciences 102, 10652–10657, doi:10.1073/pnas.0504887102 (2005).

18 Gilbert, L. A. et al. Genome-Scale CRISPR-Mediated Control of Gene Repression and Activation. Cell 159, 647–661, doi:10.1016/j.cell.2014.09.029 (2014).

19 Weltner, J. et al. Human pluripotent reprogramming with CRISPR activators. Nat Commun 9, 2643, doi:10.1038/s41467-018-05067-x (2018).

20 Diehl, W. E. et al. Ebola Virus Glycoprotein with Increased Infectivity Dominated the 2013-2016 Epidemic. Cell 167, 1088–1098 e1086, doi:10.1016/j.cell.2016.10.014 (2016).

21 Morimoto, K., Hooper, D. C., Spitsin, S., Koprowski, H. & Dietzschold, B. Pathogenicity of different rabies virus variants inversely correlates with apoptosis and rabies virus glycoprotein expression in infected primary neuron cultures. J Virol 73, 510–518, doi:10.1128/JVI.73.1.510-518.1999 (1999).

22 Duverge, A. & Negroni, M. Pseudotyping Lentiviral Vectors: When the Clothes Make the Virus. Viruses 12, doi:10.3390/v12111311 (2020).

23 Baize, S. et al. Emergence of Zaire Ebola virus disease in Guinea. N Engl J Med 371, 1418–1425, doi:10.1056/NEJMoa1404505 (2014).

24 Morimoto, K. et al. Rabies virus quasispecies: implications for pathogenesis. Proc Natl Acad Sci U S A 95, 3152–3156, doi:10.1073/pnas.95.6.3152 (1998).

25 Chan, S. Y., Speck, R. F., Ma, M. C. & Goldsmith, M. A. Distinct mechanisms of entry by envelope glycoproteins of Marburg and Ebola (Zaire) viruses. J Virol 74, 4933–4937, doi:10.1128/jvi.74.10.4933-4937.2000 (2000).

26 Hofmann-Winkler, H., Gnirss, K., Wrensch, F. & Pohlmann, S. Comparative Analysis of Host Cell Entry of Ebola Virus From Sierra Leone, 2014, and Zaire, 1976. J Infect Dis 212 Suppl 2, S172–180, doi:10.1093/infdis/jiv101 (2015).

27 Zapatero-Belinchon, F. J. et al. Characterization of the Filovirus-Resistant Cell Line SH-SY5Y Reveals Redundant Role of Cell Surface Entry Factors. Viruses 11, doi:10.3390/v11030275 (2019).

28 Iwamoto, M., Bjorklund, T., Lundberg, C., Kirik, D. & Wandless, T. J. A general chemical method to regulate protein stability in the mammalian central nervous system. Chem Biol 17, 981–988, doi:10.1016/j.chembiol.2010.07.009 (2010).

29 Horlbeck, M. A. et al. Compact and highly active next-generation libraries for CRISPR-mediated gene repression and activation. eLife 5, 914, doi:10.7554/eLife.19760 (2016).

30 Alvarez, C. P. et al. C-type lectins DC-SIGN and L-SIGN mediate cellular entry by Ebola virus in cis and in trans. J Virol 76, 6841–6844, doi:10.1128/jvi.76.13.6841-6844.2002 (2002).

31 Marzi, A. et al. Analysis of the interaction of Ebola virus glycoprotein with DC-SIGN (dendritic cell-specific intercellular adhesion molecule 3-grabbing nonintegrin) and its homologue DC-SIGNR. J Infect Dis 196 Suppl 2, S237–246, doi:10.1086/520607 (2007).

32 Simmons, G. et al. DC-SIGN and DC-SIGNR bind ebola glycoproteins and enhance infection of macrophages and endothelial cells. Virology 305, 115–123, doi:10.1006/viro.2002.1730 (2003).

33 Tuffereau, C., Benejean, J., Blondel, D., Kieffer, B. & Flamand, A. Low-affinity nerve-growth factor receptor (P75NTR) can serve as a receptor for rabies virus. EMBO J 17, 7250–7259, doi:10.1093/emboj/17.24.7250 (1998).

34 Sissoëff, L., Mousli, M., England, P. & Tuffereau, C. Stable trimerization of recombinant rabies virus glycoprotein ectodomain is required for interaction with the p75NTR receptor. J Gen Virol 86, 2543–2552, doi:10.1099/vir.0.81063-0 (2005).

35 Langevin, C., Jaaro, H., Bressanelli, S., Fainzilber, M. & Tuffereau, C. Rabies virus glycoprotein (RVG) is a trimeric ligand for the N-terminal cysteine-rich domain of the mammalian p75 neurotrophin receptor. J Biol Chem 277, 37655–37662, doi:10.1074/jbc.M201374200 (2002).

36 Southard, K. M. et al. Comprehensive transcription factor perturbations recapitulate fibroblast transcriptional states. Nat Genet 57, 2323–2334, doi:10.1038/s41588-025-02284-1 (2025).

37 Hoenen, T., Groseth, A., Callison, J., Takada, A. & Feldmann, H. A novel Ebola virus expressing luciferase allows for rapid and quantitative testing of antivirals. Antiviral Res 99, 207–213, doi:10.1016/j.antiviral.2013.05.017 (2013).

38 Kainulainen, M. H. et al. Recombinant Sudan virus and evaluation of humoral cross-reactivity between Ebola and Sudan virus glycoproteins after infection or rVSV-DeltaG-ZEBOV-GP vaccination. Emerg Microbes Infect, 2265660, doi:10.1080/22221751.2023.2265660 (2023).

39 Shalem, O. et al. Genome-scale CRISPR-Cas9 knockout screening in human cells. Science (New York, NY) 343, 84–87 (2014).

40 Biering, S. B. et al. Genome-wide bidirectional CRISPR screens identify mucins as host factors modulating SARS-CoV-2 infection. Nat Genet 54, 1078–1089, doi:10.1038/s41588-022-01131-x (2022).

41 Rebendenne, A. et al. Bidirectional genome-wide CRISPR screens reveal host factors regulating SARS-CoV-2, MERS-CoV and seasonal HCoVs. Nat Genet 54, 1090–1102, doi:10.1038/s41588-022-01110-2 (2022).

42 Zhu, S. et al. Genome-wide CRISPR activation screen identifies candidate receptors for SARS-CoV-2 entry. Sci China Life Sci 65, 701–717, doi:10.1007/s11427-021-1990-5 (2022).

43 Dinesh, R. K. et al. Membrane-wide screening identifies potential tissue-specific determinants of SARS-CoV-2 tropism. PLoS Pathog 21, e1013157, doi:10.1371/journal.ppat.1013157 (2025).

44 Liang, Z. et al. Cytotoxicity of activator expression in CRISPR-based transcriptional activation systems. Nat Commun 16, 8071, doi:10.1038/s41467-025-63570-4 (2025).

45 Wu, Q. et al. Massively parallel characterization of CRISPR activator efficacy in human induced pluripotent stem cells and neurons. Mol Cell 83, 1125–1139 e1128, doi:10.1016/j.molcel.2023.02.011 (2023).

46 Kampmann, M. CRISPRi and CRISPRa Screens in Mammalian Cells for Precision Biology and Medicine. ACS Chem Biol 13, 406–416, doi:10.1021/acschembio.7b00657 (2018).

47 Biedenkopf, N. et al. Renaming of genera Ebolavirus and Marburgvirus to Orthoebolavirus and Orthomarburgvirus, respectively, and introduction of binomial species names within family Filoviridae. Arch Virol 168, 220, doi:10.1007/s00705-023-05834-2 (2023).

48 Rhodes, A. J. Strains of rabies virus available for preparation of sylvatic rabies vaccines with special reference to vaccines prepared in cell culture. Can Vet J 22, 262–266 (1981).

49 Mentis, G. Z. et al. Transduction of motor neurons and muscle fibers by intramuscular injection of HIV-1-based vectors pseudotyped with select rabies virus glycoproteins. J Neurosci Methods 157, 208–217, doi:10.1016/j.jneumeth.2006.04.011 (2006).

50 Goldstein, A. S. et al. Purification and direct transformation of epithelial progenitor cells from primary human prostate. Nat Protoc 6, 656–667, doi:10.1038/nprot.2011.317 (2011).

51 Dull, T. et al. A third-generation lentivirus vector with a conditional packaging system. J Virol 72, 8463–8471, doi:10.1128/JVI.72.11.8463-8471.1998 (1998).

52 Ang, L. T. et al. Generating human artery and vein cells from pluripotent stem cells highlights the arterial tropism of Nipah and Hendra viruses. Cell 185, 2523–2541 e2530, doi:10.1016/j.cell.2022.05.024 (2022).

53 CDC & NIH. Biosafety in Microbiological and Biomedical Laboratories—6th Edition. (U.S. Department of Health and Human Services, 2020).

